# Comparison of RNA Isolation Methods on RNA-Seq: Implications for Differential Expression and Meta-Analyses

**DOI:** 10.1101/728014

**Authors:** Amanda N. Scholes, Jeffrey A. Lewis

**Author notes:** Corresponding author Jeffrey A. Lewis, Department of Biological Sciences University of Arkansas, 850 W. Dickson St., SCEN 601 Fayetteville, AR 72701.

## Abstract

Technical variation across different batches of RNA-seq experiments can clearly produce spurious signals of differential expression and reduce our power to detect true differences. Thus, it is important to identify major sources of these so-called “batch effects” to eliminate them from study design. Based on the different chemistries of “classic” phenol extraction of RNA compared to common commercial RNA isolation kits, we hypothesized that specific mRNAs may be preferentially extracted depending upon method, which could masquerade as differential expression in downstream RNA-seq analyses. We tested this hypothesis and found that phenol extraction preferentially isolated membrane-associated mRNAs, thus resulting in spurious signals of differential expression. Within a self-contained experimental batch (e.g. control versus treatment), the method of RNA isolation had little effect on the ability to identify differentially expressed transcripts. However, we suggest that researchers performing meta-analyses across different experimental batches strongly consider the RNA isolation methods for each experiment.

## Introduction

The decreasing cost of massively parallel sequencing had led to an explosion of transcriptomic datasets. This large number of datasets has allowed for meta-analyses, which can be valuable due to their increase in statistical power. However, researchers performing meta-analyses on transcriptomic datasets need to be cautious in their use and be aware of so-called “batch effects,” where technical differences between experimental batches can clearly produce spurious signals of differential expression and reduce our power to detect true differences.

In some cases the sources of batch effects are known and can be avoided. Some well-known batch effects include sequencing lane effects, library construction protocol, and RNA quality [1–3]. Other sources of batch effects clearly exist but remain unknown. While batch effects can sometimes be accounted for this comes with some major caveats. If the batch effect completely confounds the experimental design, for example with different sequencing lanes being used for controls and treatments, statistically accounting for the batch effect will remove any “real” signal [4]. Even in the case where the batch effect is not a complete confounder, accounting for batch can reduce our power to detect true biological signal [5]. Thus, a better understanding of the sources of batch effects can help us to avoid them.

In this study, we examined the effects of RNA isolation method as a possible source of batch effects in RNA-seq design. It is well known that the RNA distribution within cells is not uniform. Newly synthesized pre-mRNAs are processed in the nucleus before being exported. Once exported, mRNAs are frequently trafficked to specific subcellular sites as a mechanism for spatially controlling protein synthesis. Indeed, perhaps the most widespread example of mRNA localization is that used for spatial control of protein synthesis, where mRNAs encoding secreted and membrane proteins are translated at the ER membrane allowing for proper protein localization and folding [6].

Despite the widespread acknowledgement that mRNAs are differentially localized within the cell, there has been a paucity of studies examining whether “common” RNA extraction methods are equivalent in their abilities to extract differentially localized RNA species, and whether the method of RNA isolation affects our ability to detect differentially expressed transcripts. Sultan and colleagues compared two RNA isolation methods (Qiagen RNeasy kit and guanidium-phenol (TRIzol) extraction) and two library selection schemes (poly-A enrichment and rRNA depletion) on downstream transcript abundance estimates, and found that rRNA depletion was particularly sensitive to the RNA extraction method [2]. However, their comparisons were done using only two biological replicates, and they only examined transcript abundance across technical replicates and not whether the method of extraction affects the ability to detect differential expression in the types of sample comparisons that biologists frequently care about (e.g. wild-type versus mutant or treatment versus control).

Thus, we sought to systematically examine whether three common RNA isolation methods led to differences in transcript abundance and/or our ability to detect differential expression between two experimental conditions in the form of the *Saccharomyces cerevisiae* heat shock response. The different RNA isolation methods were the classic “hot acid phenol” method, and the two most commonly-used types of kits [7]—a silica-based column kit (Qiagen RNeasy Kit) and a guanidium-phenol (TRIzol)-based kit (Zymo Research Direct-zol), hereafter referred to as the Phenol, RNeasy, and Direct-zol methods. Based on the combined chemistries of sodium dodecyl sulfate (SDS) and phenol on cellular membranes [8, 9], we hypothesized that the Phenol method would better solubilize membrane-associated mRNAs. To test this hypothesis, and whether the choice of RNA isolation method had downstream effects on our ability to detect differentially expression transcripts, we collected four biological replicates of the model yeast *Saccharomyces cerevisiae* before and after a 20-minute heat shock. Importantly, each biological sample was split into three identical technical replicates that differed only in their mode of RNA isolation. This allowed us to systematically test whether the RNA isolation method affects relative transcript abundance between technical replicates, and whether that matters for differential expression analysis.

Our analysis found a shocking number of transcripts (nearly 1/3 of the genome) that appeared “differentially” expressed when comparing the Phenol method to either Kit method, and a small number of differences when comparing the Kit methods to each other. Strikingly, transcripts over-represented by the Phenol method versus either Kit method were enriched for membrane proteins, suggesting that indeed the combination of SDS, phenol, and/or heat better extracts those species of mRNA. Importantly, there were virtually no differences when comparing differential expression for the heat shock response within samples where RNA was isolated via same method. Based on these results, we strongly recommend that meta-analyses be performed on groups of experiments with common RNA isolation methods.

## Results

### Experimental setup

To test whether RNA extraction methods impact between-sample comparisons and the power to identify differentially expressed genes, we used the well-characterized yeast heat shock response as an environmental perturbation. We collected four biological replicates for comparison. For each biological replicate, three “technical replicate” samples were collected to understand the impact of RNA extraction method. The only difference between was that each technical replicate had their RNA extracted by one of three methods: classic hot acid phenol (Phenol method), a silica-based column kit (RNeasy Method) and a guanidium-phenol (TRIzol)-based kit (Direct-zol Method) (Figure 1). RNA isolated via the Phenol method was subsequently “cleaned” with a Qiagen RNeasy Kit using the optional on-column DNase treatment, thus controlling for both DNase treatment and potential differential binding of different RNA species to the column. To minimize against batch effects other than RNA extraction method, all RNA-seq libraries were constructed on the same day using an automated robotic platform, and all libraries were multiplexed and sequenced on a single lane of an Illumina HiSeq4000 instrument.

**Figure 1.**
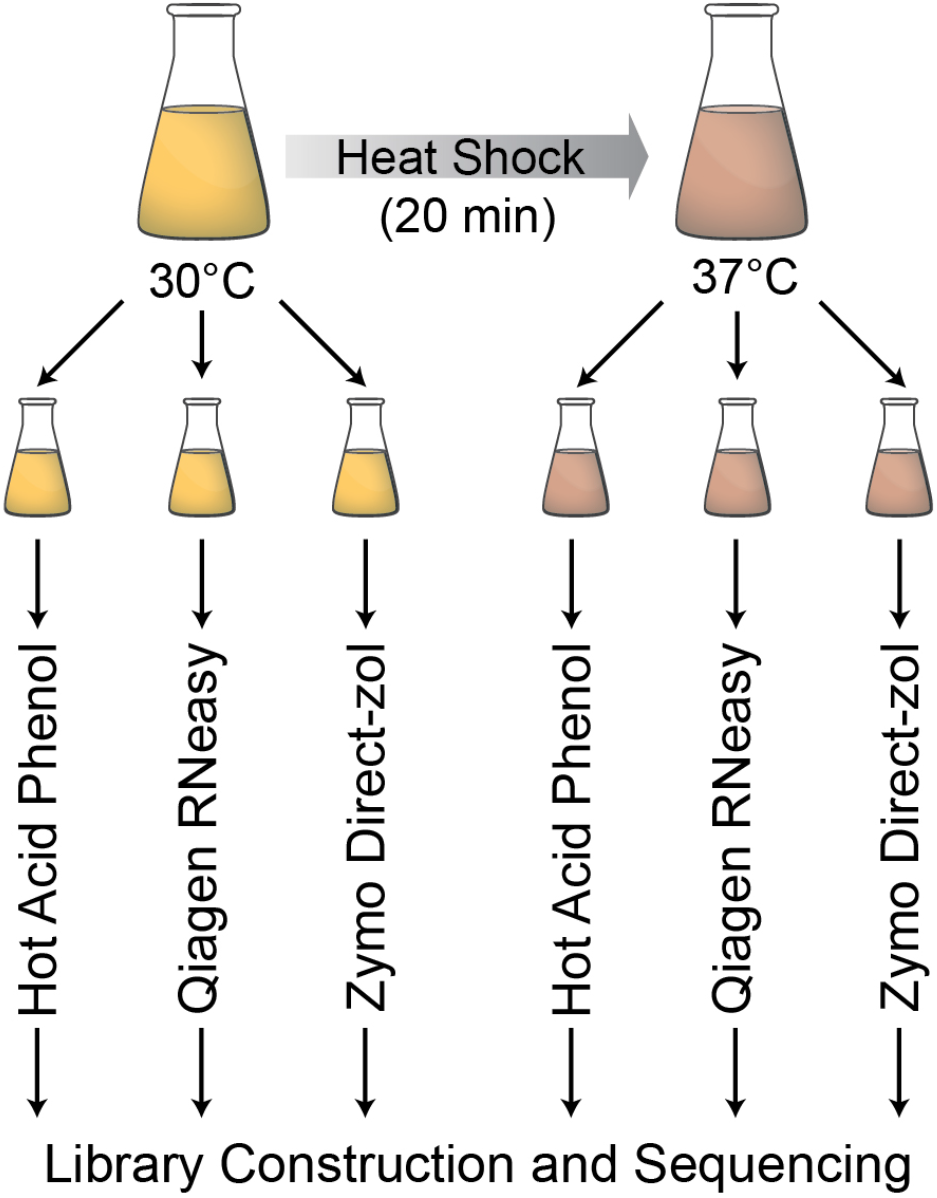
Schematic of the experimental design. Yeast cells were grown to mid-exponential phase at 30°C, unstressed control samples were collected, and then cells were shifted to a 37°C heat shock with samples collected after 20 minutes. For both unstressed and stressed cells, we collected three identical samples (technical replicates), and RNA was isolated using either hot acid phenol extraction, a Qiagen RNeasy Kit, or a Zymo Research Direct-zol RNA Kit. Libraries were constructed in a single batch using a liquid handling robot, and then were pooled and sequenced on a single Illumina HiSeq4000 lane.

### Differences in relative transcript abundance between phenol-extracted RNA and kit-extracted RNA

All of the RNA isolation methods yielded generally high quality RNA, as defined by a RIN of 9.0 or above, though the Phenol-extracted RNA averaged significantly higher RIN values than those isolated from the Direct-zol kit (9.96 vs. 9.33; *p* = 2 × 10^−6^, *t*-test) or the RNeasy kit (9.96 vs. 9.79; *p* = 0.01, *t*-test). (Table S1). The percentage of total mapped reads was similar across samples, with slight (though significant) differences (Table S2). There were larger differences in the percentage of uniquely mapped reads across RNA isolation methods (Table S2). These differences did not correlate with RNA integrity, as the Direct-zol samples had the lowest RIN values and highest uniquely and total mapped reads. Overall, we feel that the both the RNA quality and read mapping would not raise any red flags in laboratories performing RNA-seq on either their own samples, or conducting a meta-analysis, though those values can be used a factor to be controlled for in differential expression analysis [3].

We were particular interested in whether differences in the RNA isolation method could masquerade as “differential” expression due to differences in transcript quantification. We first performed principal component analysis (PCA) (Figure 2). Not surprisingly, a substantial proportion of the variance (50.5%) was explained by treatment (unstressed versus heat shock). The second principal component corresponded to RNA isolation method and explained 26.9% of the variation. Samples with RNA isolated by the two different kit methods clustered together, with the Phenol-isolated samples forming a separate cluster. It could seem counterintuitive that the Direct-zol and Phenol methods would be so dissimilar, considering that both methods use phenol. However, the Direct-zol method uses a milder detergent than SDS (sarkosyl), is performed at room temperatures instead of 65°C, and samples are exposed to phenol for 10 minutes instead of 45 minutes. We speculate that these differences, when combined with common silica-column chemistries for each kit, result in the kits behaving similarly.

**Figure 2.**
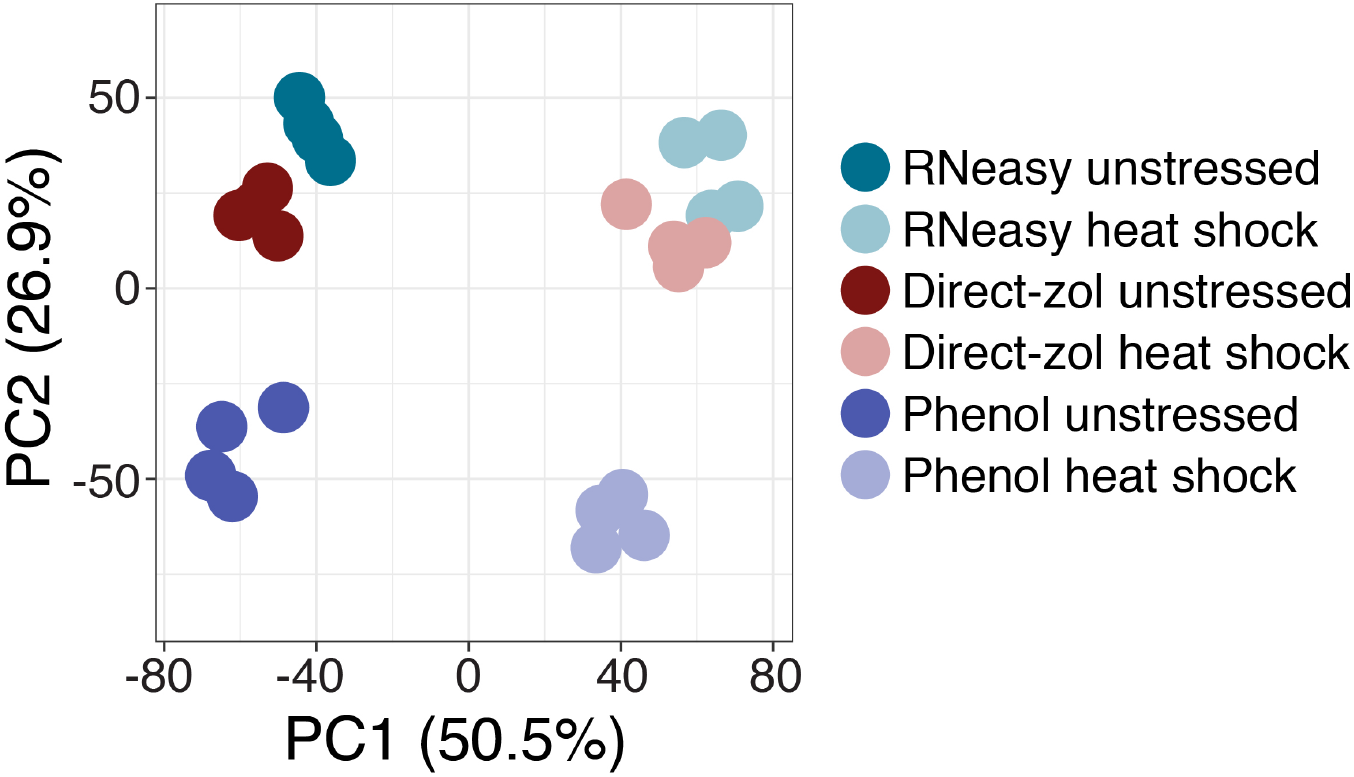
Principal component analysis (PCA) strongly implicates RNA isolation method as a batch effect. PCA on TPMs for each sample (see methods) shows clear separation on both treatment (PC1) and RNA isolation method (PC2). Kit samples were more similar to each other than they were to the phenol sample.

To visualize differences in transcript abundance across RNA isolation methods, we performed hierarchical clustering on the TPMs of the unstressed samples (Figure 3A). Hierarchical clustering of the samples largely recapitulated the patterns of PCA—again, the Phenol-isolated samples formed a discreet cluster distinct from the two kits. The RNeasy- and Direct-zol-isolated samples also had far fewer visible differences. To quantify these differences, we used edgeR to identify transcripts with significantly differential abundance in pairwise comparisons of each RNA isolation method (FDR < 0.01, see Methods). Pairwise comparisons of the Phenol method with each Kit method identified a large number of transcripts with differential abundance: 2,430 transcripts (Phenol vs. RNeasy) and 2,512 transcripts (Phenol vs. Direct-zol). Of those transcripts with differential abundance in both comparisons, 1,917 overlapped, which was highly significant (*P* = 1 × 10^−520^, Fisher’s exact test) (Figure 3C). In contrast, only 230 transcripts had differential abundance when comparing the kits to each other, suggesting only slight differences.

**Figure 3.**
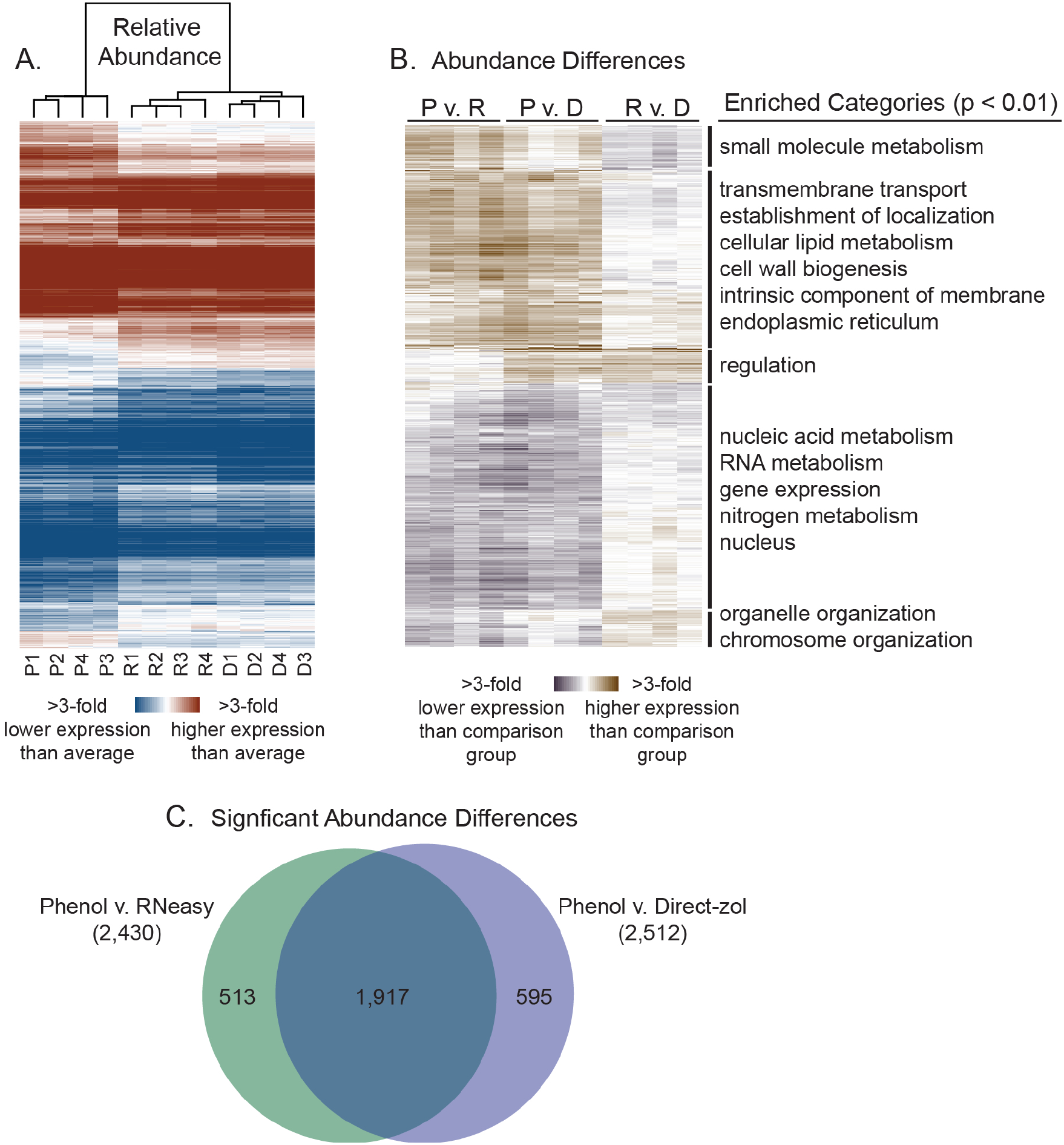
Phenol preferentially extracts mRNAs that encode for membrane proteins. (A) Hierarchical clustering of unstressed samples (P = phenol, R = RNeasy, D = Direct-zol). Clustering on relative transcript abundance (TPMs) reveals differences depending upon RNA isolation method, while clustering on sample identity shows that the phenol-isolated samples separate from both kits. Red indicates higher than average transcript abundance within a sample, and blue indicates lower than average transcript abundance. (B) Hierarchical clustering of 3,127 transcripts with significantly differential abundance (FDR < 0.01) in any pairwise comparisons between each RNA isolation method. Brown indicates higher expression than the comparison group (e.g. Phenol in the P v. R column) and violet indicates lower expression than the comparison group (e.g. RNeasy in the P v. R column). Enriched Gene Ontology (GO) categories (Bonferroni-corrected *P* < 0.01) are shown on the right. Complete GO enrichments for each cluster can be found in File S2. (C) Overlap between transcripts with significantly differential abundance (FDR < 0.01) in the Phenol v. RNeasy and Phenol v. Direct-zol comparisons.

To better visualize these differences, we performed hierarchical clustering on all 3,127 transcripts with significantly differential abundance (FDR < 0.01) in any pairwise comparison of RNA isolation method (Figure 3B). We found striking functional gene ontology (GO) enrichments for transcripts with higher or lower abundance in the phenol-extracted samples compared to both kits. Transcripts with higher abundance in phenol-extracted RNA in comparison to both kits were strongly enriched for transmembrane transport (*P* < 4 × 10^−68^), establishment of localization (*P* < 9 × 10^−54^), lipid metabolism (*P* < 1 × 10^−27^), and cell wall organization (*P* < 1 × 10^−18^). Looking more closely at the cellular component GO enrichments, transcripts with higher abundance in the phenol samples were strongly enriched for those encoding intrinsic membrane proteins (*P* < 4 × 10^−191^), as well as proteins localized to the endoplasmic reticulum (*P* < 6 × 10^−84^), cell periphery (*P* < 3 × 10^−80^), and the vacuole (*P* < 3 × 10^−53^). In contrast, mRNAs with lower relative abundance in the phenol samples were enriched for nuclear in localization (*P* < 3 × 10^−60^), and included those encoding functions related to nucleic acid metabolism (*P* < 1 × 10^−38^), RNA metabolism (*P* < 6 × 10^−28^), chromosome organization (*P* < 4 × 10^−17^), and gene expression (*P* < 8 × 10^−17^).

### Properties of transcripts with spurious differential expression

That the Phenol-isolated samples have higher transcript abundance for mRNAs encoding membrane proteins fits with the hypothesis that the Phenol method better solubilizes that species of mRNA. Another possibility is that differences in transcript degradation rates are responsible for the spurious patterns of differential expression. Because GC content and transcript length correlate with *in vivo* mRNA degradation rates [3], we examined those relationships in our data. Transcripts with significantly higher or lower abundance in Phenol-extracted samples compared to each Kit method had significantly higher GC content and gene length (Figure S1). We also examined the relationship between differential abundance and direct estimates of *in vivo* transcript stability (half-lives) from Neymotin and colleagues [10]. We did find a significant difference in the Phenol vs. Direct-zol comparison, but not for the Phenol vs. RNeasy comparison. To determine how much of the variation was explained by GC content, gene length, and transcript half-life, we performed linear regression of those parameters on the average fold changes for phenol-extracted samples vs. the kits. Both GC content and transcript length showed weak to moderate correlation (*r* = 0.06 – 0.32) with log_2_ fold changes, depending upon the comparison group, while estimated *in vivo* half-life weakly correlated with log_2_ fold changes in either comparison (Table S3). Because differences in GC content and length are associated with differences in transcript degradation rates *in vitro* [3], we repeated the edgeR analysis using RIN as a factor. We expected that because the RIN values for the Direct-zol samples were all lower than the others, it would eliminate most of the signal for differential expression. This turned out to be correct—we identified 788 “differentially” expressed genes in the Phenol vs. Direct-zol comparison compared to 2,513 when RIN was not included as a factor. The surviving differentially expressed transcripts with higher expression in the Phenol-isolated samples relative to the Direct-zol isolated samples were still strongly enriched for those encoding intrinsic membrane proteins (*P* < 3 × 10^−100^). Because the RNeasy-isolated samples had relatively high RIN values relative to the Direct-zol-isolated samples, the vast majority were retained as differentially expressed when accounting for RIN in the edgeR QL model (2,362 / 2,430). Because of the substantial overlap between genes called as differentially expressed in the Phenol vs. RNeasy and Phenol vs. Direct-zol comparisons, we hypothesize that the that differing chemistries in the extraction are responsible for the batch effect, and not RNA degradation (see Discussion).

### Differences in RNA isolation method have little effect on the ability to detect differential expression with a batch

The striking differences in transcript abundance depending on RNA isolation could conceivably affect the ability to detect differential expression. To test this, we examined our ability to detect differential expression in cells shifted from 30°C to 37°C for 20 minutes—the classic yeast heat shock response. We identified ~3,800 differentially expressed transcripts for all three RNA isolation methods, with substantial overlap for all three (Figure 4). Hierarchical clustering yielded no clear pattern among differentially expressed transcripts that were missed in one sample set over another (Figure 4). We also detected zero transcripts that had significant fold change differences in their heat shock response in any pairwise comparison between RNA isolation methods (File S2). We hypothesize that at sufficient sequencing depth, the ability to detect differential expression is robust to the modest differences in transcript counts caused by differences in RNA isolation method.

**Figure 4.**
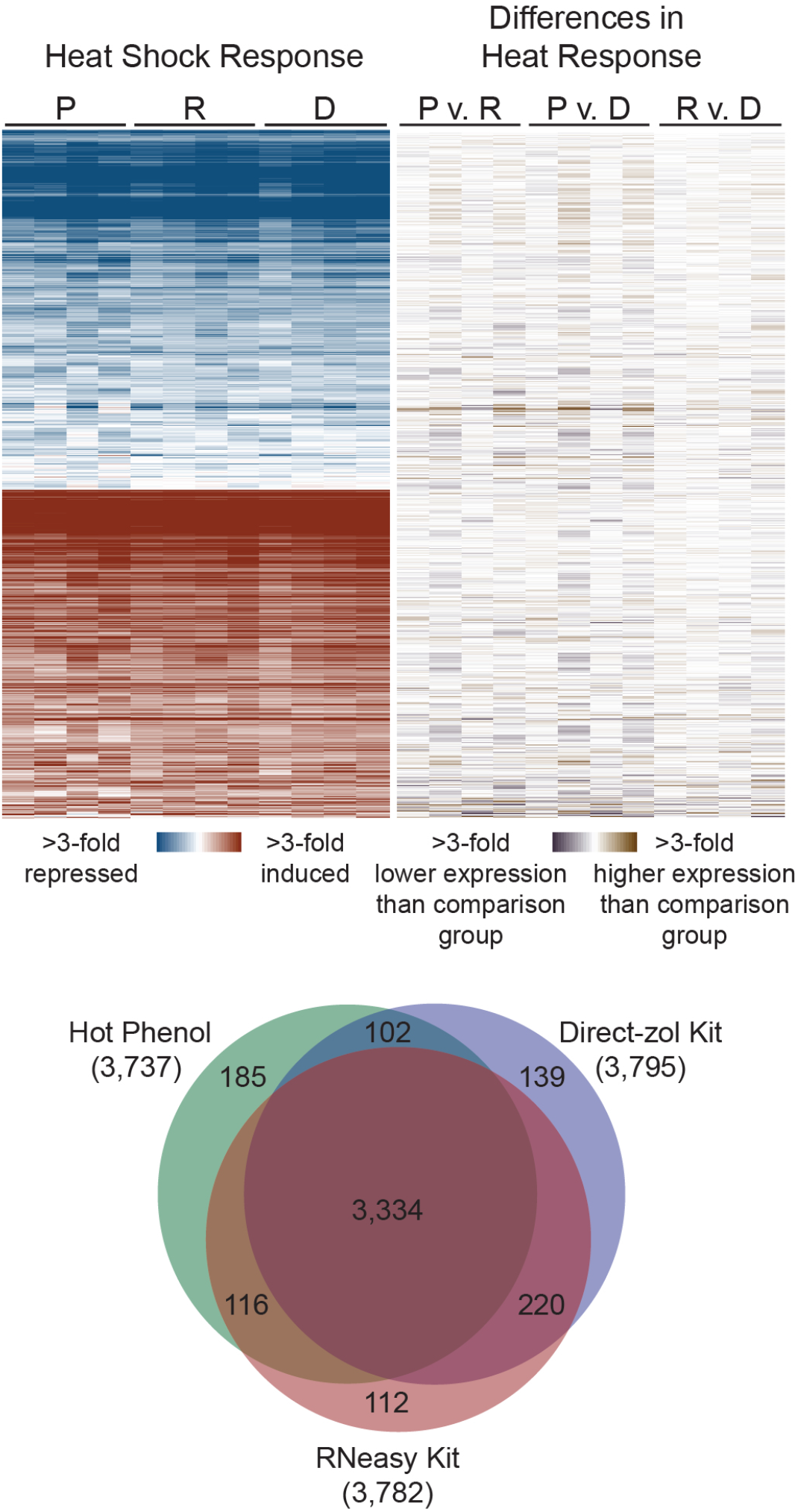
The method of RNA extraction has little effect on differential expression analysis. Hierarchical clustering of median-centered log_2_-fold TPM changes for 4,232 transcripts that were differentially expressed in response to heat (FDR < 0.01) in at least one set of samples (P = phenol, R = RNeasy, D = Direct-zol). The left portion of the heat map displays gene expression changes during heat shock across the four biological replicates, with red indicated genes induced by heat shock, and blue indicating genes repressed by heat shock. The right portion shows differences in abundance in pairwise comparisons between each RNA isolation method, with brown indicating higher expression than the comparison group, and violet indicating lower expression than the comparison group. The Venn Diagram indicates overlap between differentially expressed genes in the Phenol, RNeasy, and Direct-zol isolated samples.

## Discussion

In this study, we tested whether differences in RNA isolation method affect relative transcript abundance between samples, and whether the RNA isolation method impacts our ability to detect differential expression. Our results suggest that differences in RNA isolation method can substantially affect relative transcript abundance, and we see thousands of differences in transcript abundance when comparing hot acid phenol extraction with an RNeasy or Direct-zol kit. It is well established that mRNAs encoding membrane and secreted proteins are anchored to the membrane during translation [11]. That transcripts with higher abundance in the Phenol-isolated samples are strongly enriched for encoding membrane proteins suggests that the Phenol method better solubilizes those mRNAs. Because relatively more membrane-associated mRNAs are being extracted, there must be relatively less abundance of other mRNAs. Thus, we see decreased abundance of certain nuclear transcripts, which were already more lowly expressed, and thus likely more sensitive to appearing “repressed.”

We disfavor the alternative hypothesis that we are capturing differences in transcript degradation rates for a number of reasons. First, while we do see differences in RIN values across the different RNA isolation methods, the differences are relatively small, and our RIN values are all much higher than the points where other studies identified them as confounding for RNA-seq analysis [3, 12]. Second, it is likely that any degradation that is occurring in our samples is happening *in vitro* during RNA isolation, and Opitz and colleagues have found that *in vitro* RNA degradation rates are likely relatively equal across transcripts and thus have little effect on differential expression analysis [13]. And while RNA degradation rates *in vivo* are strongly biased and can lead to spurious functional enrichments in downstream analysis, we found little relationship between estimated mRNA half-lives from [10] and fold-changes in comparisons between kits. Only one of the Phenol vs. Kit method comparisons showed a significant difference in half-lives, but the correlation was still rather poor (*r*^2^ = 0.02). And while transcripts with higher relative abundance in the phenol-extracted samples versus the kits had higher GC content and gene length, which both correlate with higher *in vivo* degradation rates [3], the correlation between those parameters and fold-change differences is not strong (Table S3). Notably, GC content and gene length are not random, and membrane proteins tend to be longer and have higher GC content than average [14, 15]. Finally, if RNA degradation is responsible, it is somewhat hard to reconcile that we see similar patterns of “differential” expression when comparing the Phenol vs. Direct-zol and Phenol vs. RNeasy samples, even though the RNeasy samples have quite a bit higher RIN values.

Regardless of the cause of these differences between hot-phenol extracted samples and kits, it clear that this can represent a large source of batch-effect variation between samples whose RNA has been isolated via different methods. Within an individual lab, we are largely agnostic. The method of RNA isolation had little effect on the ability to identify differentially expressed transcripts in our heat shock test case. Thus, experiments within a single lab are unlikely to be affected by the choice of RNA isolation method as long as the same method is used throughout an experiment. For meta-analyses however, we recommend that researchers make every attempt to only compare experiments where the RNA isolation methods are similar.

## Materials and Methods

### Yeast Growth and Sampling Procedures

All experiments were performed using yeast strain BY4741 (S288c background; MATa *his3Δ1 leu2Δ0 met15Δ0 ura3Δ0*), obtained from Open Biosystems. To compare RNA isolation methods, we collected three identical 10-ml ‘technical’ replicates for each biological replicate (4 biological replicates in total). Cells were grown >8 generations in 100-ml synthetic complete medium (SC) [Sherman 2002] at 30°C with orbital shaking (270 rpm) shaking to mid-exponential phase (OD_600_ of 0.3 – 0.6), and 10-ml samples were removed representing the unstressed control. For heat shock treatment, one volume of 55°C medium was added to the remaining culture, immediately bringing the final temperature to 37°C, and the culture was incubated at 37°C for another 20 minutes before removing 10-ml samples. Both unstressed and heat shocked cells were collected by centrifugation at 1,500 × *g* for 3 minutes, and cell pellets were flash frozen in liquid nitrogen and stored at −80°C until processing.

### RNA Isolation Methods

#### Hot Phenol Isolation

Cells were lysed and RNA was isolated using a standard hot phenol method as described [16], and a detailed protocol can be found on the protocols.io repository under DOI dx.doi.org/10.17504/protocols.io.inwcdfe. Briefly, 1 volume of acid saturated phenol and 1 volume of lysis buffer (10 mM Tris-HCl pH 7.4, 10 mM EDTA, 0.5% SDS) were added to frozen cell pellets, vortexed, and then placed in a 65°C preheated Multi-Therm incubated vortexer (Benchmark Scientific) at 1500 rpm for 45 minutes. Samples were centrifuged for 10 min at 4°C at maximum speed in a microcentrifuge, extracted once more with phenol, once with chloroform, and then precipitated overnight at −20°C with 0.1 volumes of sodium acetate (pH 5.2) and 2.5 volumes of 100% ethanol. Precipitated RNA was washed once with 70% ethanol and then resuspended in TE (10 mM Tris-HCl pH 8.0, 1 mM EDTA). The phenol extracted RNA was then ‘cleaned’ using an RNeasy Miniprep Kit with optional on-column DNase treatment according to the manufacturer’s instructions.

### RNA Isolation with Two Different Miniprep Kits

RNA was extracted using two different kits: the Qiagen RNeasy Mini Kit (Cat. 74104) and the Zymo Research Direct-zol RNA Miniprep Kit (Cat. R2050). Cell concentrations were all below the maximum recommendation of 5 × 10^7^ cells from both manufacturers (ranging from 2.5 × 10^7^ – 4.5 × 10^7^ cells). For both kits, we mechanically lysed cells with a Beadbeater-24 (3,500 oscillations/minute, 45 seconds on ice between cycles). Mechanical lysis was performed in 2-ml screw-capped tubes containing an equal volume (600 μl) of lysis buffer (RLT for RNeasy or TRI reagent for Direct-zol) and acid-washed glass beads (425-600 micron, Sigma-Aldrich). RNA was then purified according to each manufacturer’s protocol for yeast, including the optional on-column DNase digestion. For all samples, RNA was quantitated using a Qubit RNA HS Assay kit and Qubit fluorometer according to the manufacturer’s instructions. The RNA integrity number (RIN) for each sample was measured using an Agilent 2200 TapeStation. RNA concentrations and RIN values for each sample can be found in Table S1.

### RNA Sequencing and Analysis

RNA-seq libraries were prepared from polyA-enriched RNA using the KAPA Biosystems mRNA HyperPrep Kit (KK8581) and KAPA Single-Indexed Adapter Set A+B (KK8700), according to manufacturer’s instructions. We started with 500 ng total RNA, fragmentation time (6 min) was optimized to generate 200-300-nt RNA fragments, and the libraries were amplified with 9 cycles of PCR. All libraries were constructed in a single batch through an automated Eppendorf epMotion 5075 liquid handling robot, and detailed a protocol can be found on protocols.io under DOI dx.doi.org/10.17504/protocols.io.uueewte. cDNA libraries were sequenced on a HiSeq4000 at the University of Chicago Genomics Facility, generating single-end 50-bp reads.

Reads were trimmed of low-quality reads and adapter sequence (KAPA v1 indices) using Trimmomatic (version 0.32) [17], with the following commands: ILLUMINACLIP:Kapa_indices.fa:2:30:10 LEADING:3 TRAILING:3 MAXINFO:40:0.4 MINLEN:40 . Reads were mapped to the S288c genome (version Scer3), using STAR (version 020201) [18]. Mapping statistics can be found in Table S2. Transcripts per million (TPM) and expected counts for each gene were calculated using RSEM (version 1.3.1) [19]. The RSEM output can be found in File S2.

Differential expression analysis was conducted using the Bioconductor package edgeR (version 3.22.3) using the quasi-likelihood (QL) framework. For the QL model, sample type (i.e. Phenol unstressed, Phenol heat shock, RNeasy unstressed…) and biological replicate were used as factors. To account for differences in RIN across samples, we also performed a separate analysis that included sample type, replicate, and RIN as factors in the model. To control for differences in sequencing depth across samples, the edgeR function thincounts was used to randomly subsample counts across all samples to be equal to the sample with the lowest number of total counts (8,678,188). Only genes with at least 1 count per million (CPM) in at least one condition were included in analyses All RNA-seq data are available through the National Institutes of Health Gene Expression Omnibus (GEO) database under accession no. GSE135430, and the edgeR outputs can be found in File S2.

Principle component analysis (PCA) was performed using ClustVis [20] on ln-transformed TPM values for all transcripts included in the differential expression analysis, using unit variance scaling and singular value decomposition. Hierarchical clustering was performed with Cluster 3.0 (http://bonsai.hgc.jp/~mdehoon/software/cluster/software.htm) using uncentered Pearson correlation and centroid linkage as the metric [21]. RNA-seq samples were weighted using a cutoff value of 0.4 and an exponent value of 1. Functional enrichments of gene ontology (GO) categories were performed using GO-TermFinder (https://go.princeton.edu/cgi-bin/GOTermFinder)[22], with Bonferroni-corrected *P*-values < 0.01 taken as significant. Complete lists of enriched categories can be found in File S3.

## Acknowledgements

We thank Zymo Research for providing a free sample of the Direct-zol RNA Kit, Dr. Elizabeth Ruck for running samples on the TapeStation, and Dr. Andrew Alverson for the use of equipment. This work was supported in part by National Science Foundation Grant No. IOS-1656602 (JAL), the Arkansas Biosciences Institute (Arkansas Settlement Proceeds Act of 2000) (JAL), and a Research Assistantship provided through the University of Arkansas Cell and Molecular Biology Graduate Program (ANS). The funders had no role in study design, data collection and interpretation, or the decision to submit the work for publication.

---

**Figure S1.**
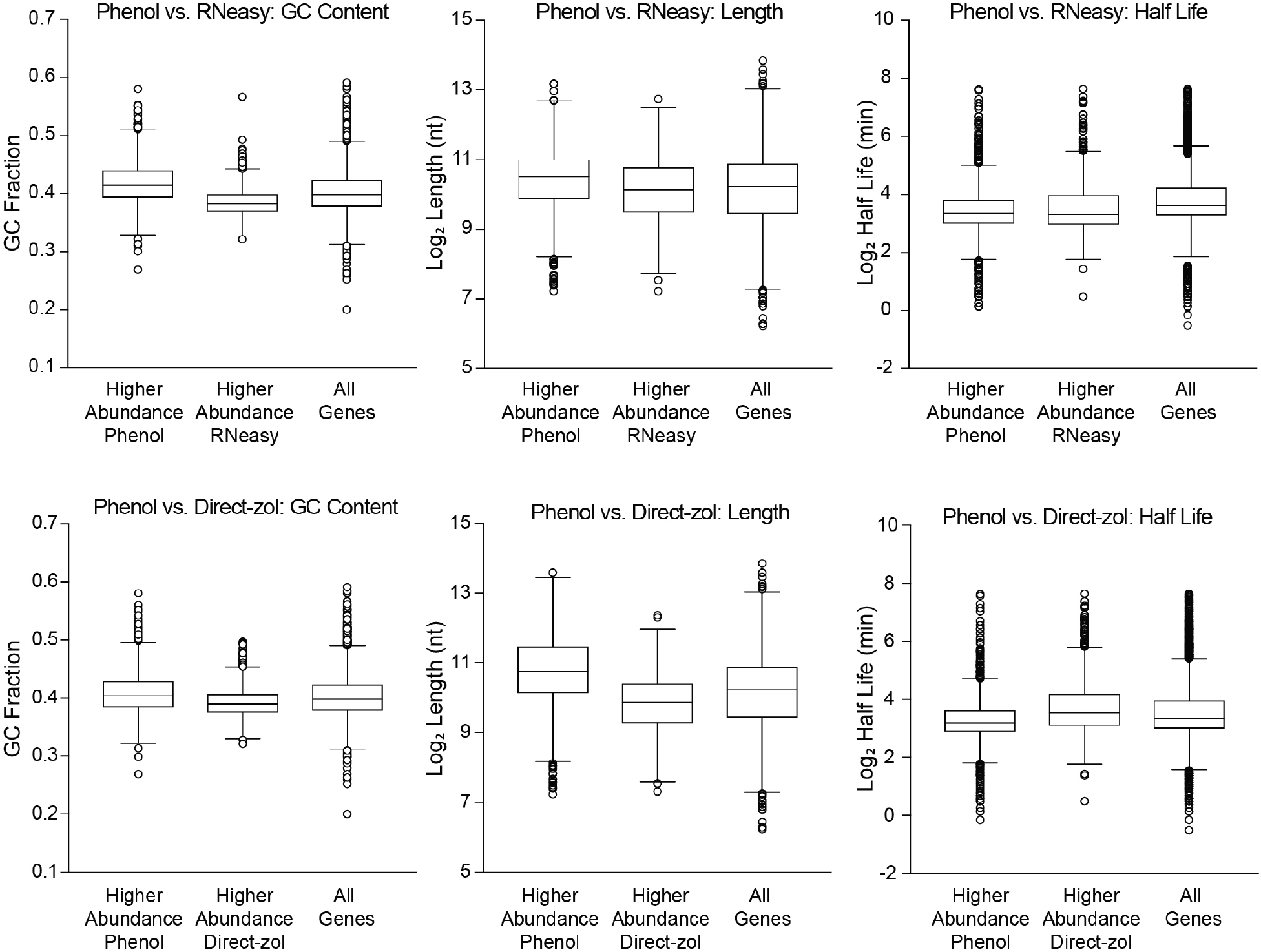
Properties of transcripts with differential abundance depending upon RNA isolation method. Boxplots depicting the GC content (left), transcript length (middle), and *in vivo* half-life (obtained from [10]) for all genes with significantly higher abundance relative to their comparison group (FDR < 0.01), and all genes that passed filtering thresholds for differential expression analysis. For the Phenol vs. RNeasy comparisons, mean differences were significant by Mann Whitney *U* test for GC content (*P* < 1 × 10^−15^) and length (*P* < 1 × 10^−15^). For the Phenol vs. Direct-zol comparisons, mean differences were significant by Mann Whitney *U* test for GC content (*P* < 1 × 10^−15^), length (*P* < 1 × 10^−15^), and half-life (*P* < 1 × 10^−15^).

**Table S1.** RNA concentrations and integrity (RIN) scores.

| Sample ID | Concentration<br>(ng / μl) | RIN |
| --- | --- | --- |
| Phenol unstressed Rep1 | 1,100 | 9.9 |
| Phenol unstressed Rep2 | 1,080 | 9.9 |
| Phenol unstressed Rep3 | 960 | 9.9 |
| Phenol unstressed Rep4 | 720 | 10 |
| Phenol heat shock Rep1 | 610 | 10 |
| Phenol heat shock Rep2 | 710 | 10 |
| Phenol heat shock Rep3 | 770 | 10 |
| Phenol heat shock Rep4 | 610 | 10 |
| RNeasy unstressed Rep1 | 580 | 9.8 |
| RNeasy unstressed Rep2 | 470 | 10 |
| RNeasy unstressed Rep3 | 530 | 9.9 |
| RNeasy unstressed Rep4 | 460 | 9.7 |
| RNeasy heat shock Rep1 | 400 | 9.7 |
| RNeasy heat shock Rep2 | 460 | 9.6 |
| RNeasy heat shock Rep3 | 430 | 9.8 |
| RNeasy heat shock Rep4 | 360 | 9.8 |
| Direct-zol unstressed Rep1 | 590 | 9.4 |
| Direct-zol unstressed Rep2 | 470 | 9.4 |
| Direct-zol unstressed Rep3 | 620 | 9.5 |
| Direct-zol unstressed Rep4 | 330 | 9.3 |
| Direct-zol heat shock Rep1 | 260 | 9.0 |
| Direct-zol heat shock Rep2 | 260 | 9.4 |
| Direct-zol heat shock Rep3 | 560 | 9.3 |
| Direct-zol heat shock Rep4 | 320 | 9.4 |

**Table S2.** Summary of mapping statistics.

| Sample ID | Input Reads<sup>1</sup> | % Uniquely Mapped | % Mapped |
| --- | --- | --- | --- |
| Phenol unstressed Rep1 | 13,270,392 | 83.58 | 96.21 |
| Phenol unstressed Rep2 | 12,695,531 | 83.88 | 95.76 |
| Phenol unstressed Rep3 | 12,856,634 | 85.13 | 96.32 |
| Phenol unstressed Rep4 | 15,327,828 | 84.20 | 95.97 |
| Phenol heat shock Rep1 | 10,900,974 | 86.5 | 96.57 |
| Phenol heat shock Rep2 | 13,452,555 | 86.47 | 96.83 |
| Phenol heat shock Rep3 | 14,946,326 | 87.32 | 96.82 |
| Phenol heat shock Rep4 | 19,514,083 | 87.19 | 96.89 |
| RNeasy unstressed Rep1 | 16,773,488 | 78.47 | 95.77 |
| RNeasy unstressed Rep2 | 18,735,607 | 81.06 | 95.89 |
| RNeasy unstressed Rep3 | 13,006,805 | 81.41 | 95.57 |
| RNeasy unstressed Rep4 | 20,186,658 | 80.53 | 95.38 |
| RNeasy heat shock Rep1 | 12,162,093 | 81.63 | 95.76 |
| RNeasy heat shock Rep2 | 13,405,344 | 82.18 | 96.3 |
| RNeasy heat shock Rep3 | 16,331,674 | 83.91 | 95.62 |
| RNeasy heat shock Rep4 | 22,683,053 | 80.05 | 95.87 |
| Direct-zol unstressed Rep1 | 16,815,144 | 86.38 | 96.91 |
| Direct-zol unstressed Rep2 | 17,016,270 | 87.04 | 96.95 |
| Direct-zol unstressed Rep3 | 11,598,964 | 85.87 | 96.73 |
| Direct-zol unstressed Rep4 | 14,150,816 | 87.43 | 96.83 |
| Direct-zol heat shock Rep1 | 14,678,584 | 88.02 | 96.72 |
| Direct-zol heat shock Rep2 | 16,124,462 | 88.79 | 97.27 |
| Direct-zol heat shock Rep3 | 16,753,180 | 87.92 | 96.89 |
| Direct-zol heat shock Rep4 | 21,019,701 | 87.75 | 97.14 |

<sup>1</sup> Only reads surviving trimming were used as the input for STAR mapping.

**Table S3.**
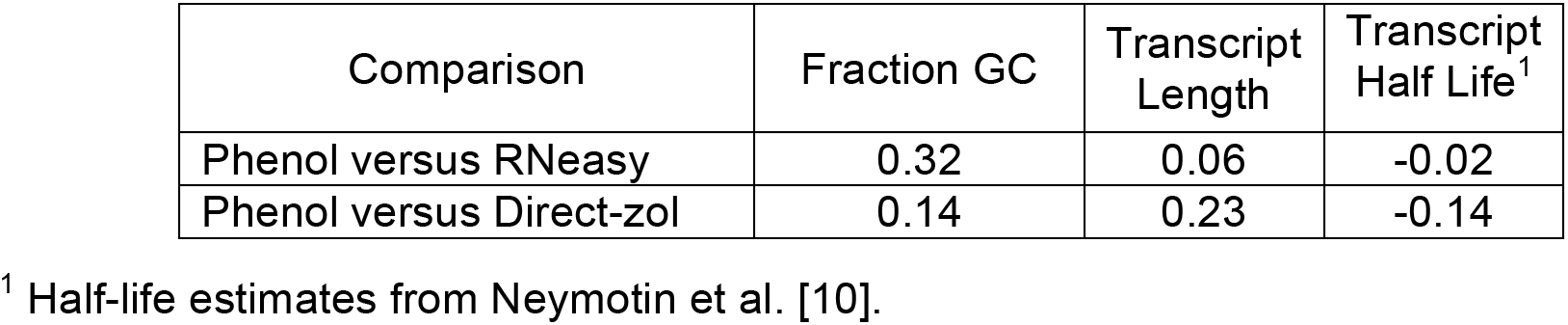
Correlation coefficient (***r***) for log_2_ fold-changes versus each factor.

## References

1. Li S, Labaj PP, Zumbo P, Sykacek P, Shi W, Shi L, et al. Detecting and correcting systematic variation in large-scale RNA sequencing data. Nat Biotechnol. 2014;32(9):888–895. doi: 10.1038/nbt.3000 PMID: 25150837

2. Sultan M, Amstislavskiy V, Risch T, Schuette M, Dokel S, Ralser M, et al. Influence of RNA extraction methods and library selection schemes on RNA-seq data. BMC Genomics. 2014;15(675. doi: 10.1186/1471-2164-15-675 PMID: 25113896

3. Gallego Romero I, Pai AA, Tung J, Gilad Y. RNA-seq: impact of RNA degradation on transcript quantification. BMC Biol. 2014;12(42. doi: 10.1186/1741-7007-12-42 PMID: 24885439

4. Gilad Y, Mizrahi-Man O. A reanalysis of mouse ENCODE comparative gene expression data. F1000Res. 2015;4(121. doi: 10.12688/f1000research.6536.1 PMID: 26236466

5. Goh WWB, Wang W, Wong L. Why Batch Effects Matter in Omics Data, and How to Avoid Them. Trends Biotechnol. 2017;35(6):498–507. doi: 10.1016/j.tibtech.2017.02.012 PMID: 28351613

6. Lerner RS, Seiser RM, Zheng T, Lager PJ, Reedy MC, Keene JD, et al. Partitioning and translation of mRNAs encoding soluble proteins on membrane-bound ribosomes. RNA. 2003;9(9):1123–1237. doi: 10.1261/rna.5610403 PMID: 12923260

7. Johnson M. Kits for RNA Extraction, Isolation, and Purification. Materials and Methods. 2012;2(201. doi: 10.13070/mm.en.2.201 PMID:

8. Kragh-Hansen U, le Maire M, Moller JV. The mechanism of detergent solubilization of liposomes and protein-containing membranes. Biophys J. 1998;75(6):2932–2946. doi: 10.1016/S0006-3495(98)77735-5 PMID: 9826614

9. Davidson PM, Branden AL. Antimicrobial Activity of Non-Halogenated Phenolic Compounds. J Food Prot. 1981;44(8):623–632. doi: 10.4315/0362-028X-44.8.623 PMID: 30836539

10. Neymotin B, Athanasiadou R, Gresham D. Determination of in vivo RNA kinetics using RATE-seq. RNA. 2014;20(10):1645–1652. doi: 10.1261/rna.045104.114 PMID: 25161313

11. Chartron JW, Hunt KC, Frydman J. Cotranslational signal-independent SRP preloading during membrane targeting. Nature. 2016;536(7615):224–228. doi: 10.1038/nature19309 PMID: 27487213

12. Chen EA, Souaiaia T, Herstein JS, Evgrafov OV, Spitsyna VN, Rebolini DF, et al. Effect of RNA integrity on uniquely mapped reads in RNA-Seq. BMC Res Notes. 2014;7(753. doi: 10.1186/1756-0500-7-753 PMID: 25339126

13. Opitz L, Salinas-Riester G, Grade M, Jung K, Jo P, Emons G, et al. Impact of RNA degradation on gene expression profiling. BMC Med Genomics. 2010;3(36. doi: 10.1186/1755-8794-3-36 PMID: 20696062

14. D’Onofrio G, Jabbari K, Musto H, Bernardi G. The correlation of protein hydropathy with the base composition of coding sequences. Gene. 1999;238(1):3–14. doi: 10.1016/s0378-1119(99)00257-7 PMID: 10570978

15. Young MD, Wakefield MJ, Smyth GK, Oshlack A. Gene ontology analysis for RNA-seq: accounting for selection bias. Genome Biol. 2010;11(2):R14. doi: 10.1186/gb-2010-11-2-r14 PMID: 20132535

16. Gasch AP. Yeast genomic expression studies using DNA microarrays. Methods Enzymol. 2002;350(393-414. doi: PMID: 12073326

17. Bolger AM, Lohse M, Usadel B. Trimmomatic: a flexible trimmer for Illumina sequence data. Bioinformatics. 2014;30(15):2114–2120. doi: 10.1093/bioinformatics/btu170 PMID: 24695404

18. Dobin A, Davis CA, Schlesinger F, Drenkow J, Zaleski C, Jha S, et al. STAR: ultrafast universal RNA-seq aligner. Bioinformatics. 2013;29(1):15–21. doi: 10.1093/bioinformatics/bts635 PMID: 23104886

19. Li B, Dewey CN. RSEM: accurate transcript quantification from RNA-Seq data with or without a reference genome. BMC Bioinformatics. 2011;12(323. doi: 10.1186/1471-2105-12-323 PMID: 21816040

20. Metsalu T, Vilo J. ClustVis: a web tool for visualizing clustering of multivariate data using Principal Component Analysis and heatmap. Nucleic Acids Res. 2015;43(W1):W566–W570. doi: 10.1093/nar/gkv468 PMID: 25969447

21. Eisen MB, Spellman PT, Brown PO, Botstein D. Cluster analysis and display of genome-wide expression patterns. Proc Natl Acad Sci U S A. 1998;95(14863–14868. doi: PMID: 9843981

22. Boyle EI, Weng S, Gollub J, Jin H, Botstein D, Cherry JM, et al. GO∷TermFinder--open source software for accessing Gene Ontology information and finding significantly enriched Gene Ontology terms associated with a list of genes. Bioinformatics. 2004;20(18):3710–3715. doi: 10.1093/bioinformatics/bth456 PMID: 15297299

